# Serotonergic circuit architecture underlies sex dimorphism in anxiogenic states

**DOI:** 10.64898/2026.09.15.751692

**Authors:** Noorya Yasmin Ahmed, Yillcer Molina, Alvaro Ballesteros Gonzalez, Alicia Juan, William Kwan, Anna Nguyen, Candela Barettino, Ines Botia, Ilvana Ziko, Dhanisha J Jhaveri, Isabel Del Pino, Nathalie Dehorter

## Abstract

The serotonergic system underpins anxiety-like behavior vital for the adaptive response to a new environment. However, our understanding of the neural circuitry underlying the diversity of anxiety-like behaviors across individuals is limited. Whilst previous studies have characterized diverse features related to the neurochemical substrate of serotonergic neurotransmission and behavioral presentation between males and females, no investigations have fully addressed the sexual dimorphism in the structural and functional connectivity of serotonergic circuits. Here, using functional FosTRAP connectivity analysis, we found sex-dependent functional differences in the serotonergic neurons of the dorsal raphe nucleus (DRN), such as enriched connectivity to the amygdala in females, compared to male mice. We also discovered previously unidentified morpho-functional sex-specific differences in DRN serotonergic circuitry that reflects the disparity in behavior presentation. To understand the molecular basis of serotonergic circuit architecture differences between sexes, we leveraged on single-cell RNA sequencing dataset from DRN serotonergic cells taking sex as a biological variable and spotted some gene candidates involved in neural circuit wiring. Altering the expression of the receptor tyrosine kinase Erbb4 in serotonergic circuits, altered the axonal connectivity to downstream targets and shifted sex-specific behavior. Overall, this study provides an essential foundation to delineate the molecular basis and connectivity pattern mechanisms underlying the serotonergic system-dependent sex-specific anxiety behavior. Since alterations in serotonin function play a vital role in adaptive behavior, as well as various pathologies including chronic anxiety and depression, this study may advance our understanding of sex-biases in presentation and treatment response of neuropsychiatric disorders.

**Breakthroughs of this work:** We identified structural sex-specific features of serotonergic innervation from the DRN and revealed molecular determinants of sex-dimorphic serotonergic connectivity.

**Teaser:** Sex dimorphism in anxiety is shaped by distinct connectivity patterns and molecular programs within serotonergic circuits in mice.

## INTRODUCTION

Anxiety is a fundamental adaptive state that enables organisms to detect potential threats and adjust behaviour accordingly. By promoting appropriate avoidance, vigilance and risk assessment, anxiety helps individuals navigate uncertain or threatening environments and is therefore tightly linked to adaptive behavioural responses. When these responses become dysregulated, however, anxiety can impair rather than support adaptive behaviour (*1*, *2*). Understanding the neural mechanisms that regulate anxiety is therefore essential for understanding how behavioural adaptation emerges and varies across individuals.

The serotonergic system is a key regulator of anxiety and adaptive responses to environmental challenges. The dorsal raphe nucleus (DRN), which contains most forebrain-projecting serotonergic neurons, provides extensive innervation to brain regions involved in emotional and affective processing, including the amygdala and hippocampus. This serotonergic network is therefore well positioned to coordinate the response of different brain regions concurrently in response to environmental information, therefore modulating the behavioural responses to anxiogenic stimuli(*3–5*). Importantly, its development and function are shaped by biological sex (*6*), providing a potential substrate for sex-specific differences in adaptive and anxiety-related behaviours.

Historically, sex-dependent variation in serotonin signalling has been attributed to differences in receptor expression, transporter abundance, and neurotransmitter synthesis in both rodents (*7*) and humans (*8*, *9*). For example, the human brain exhibits sexually dimorphic expression of the serotonin transporter (SERT) and the 5-HT1A receptor (*10*), as well as differences in serotonin synthesis within key limbic regions such as the amygdala and hippocampus (*9*). Given the well-established regulation of serotonin signalling by gonadal hormones, including testosterone, estrogens, and progesterone–(*7*, *11*, *12*), it has generally been assumed that sex-dependent behavioral and physiological differences arise primarily through hormone-mediated modulation of neurotransmission via differences in the neurochemical substrate (serotonergic receptors, transporters) and serotonin load in the adult brain.

Although considerable attention has been devoted to understanding how extrinsic factors, particularly sex hormones, influence brain function and behavior, comparatively little is known about the intrinsic molecular mechanisms that drive the emergence of sex-specific neural circuit function and to whether sex-specific function it rooted on neural circuit architecture differences. Addressing this gap is particularly important because many neuropsychiatric and neurodevelopmental disorders exhibit marked sex biases in prevalence, developmental trajectory, and symptom presentation. For example, anxiety and mood disorders occur more frequently in females (*13*), whereas autism spectrum disorder is diagnosed substantially more often in males (*14*). These observations suggest that sex-specific developmental programs contribute to the organization of neural circuits underlying behavior and highlight the need to identify the molecular mechanisms through which biological sex shapes brain connectivity and function.

In the present study, we investigated whether anxiety-like behavior in mice, particularly their response to a novel, anxiogenic environment, is sex-dependent and whether sex-differences in behavior extend beyond neurochemical regulation to the structural organization of serotonergic circuits. By systematically mapping serotonergic projections throughout the forebrain, we uncovered sex-biased patterns of axonal innervation within regions implicated in emotional processing, including circuits involved in anxiety-related behaviors. These findings identify anatomical organization of the serotonergic system as a previously underappreciated substrate of sex-specific behavioral adaptation. Because axonal connectivity emerges through tightly regulated molecular programs, we next sought to identify molecular mechanisms that contribute to the establishment of these sexually divergent circuit architectures. Through interrogation of transcriptomic datasets, we identified the receptor tyrosine kinase ErbB4 as a candidate regulator of serotonergic circuit development. ErbB4 exhibited sex-biased expression at both the transcript and protein levels in serotonergic neurons. Finally, selective manipulation of ErbB4 expression in serotonergic cells altered their projection patterns and the behavior in sex-dependent manner, demonstrating a causal role for this pathway in shaping sexually serotonergic connectivity that underlies sex-dimorphisms in anxiety-like behavior. Together, our findings reveal a developmental mechanism underlying sex-specific organization of serotonergic circuits and establish structural circuit architecture as a key contributor to behavioral adaptation differences between males and females.

## MATERIAL AND METHODS

### Mice

We generated *Pet-Cre^+/−^; RCE^f/+^* mice (postnatal day (P) 60-70) at the Australian Phenomics Facility, Australian National University (APF, ANU) from crosses with *Pet-Cre^+/−^* mice (generously gifted by Professor Greg Stuart, ANU) and *RCE^f/f^*mice (generously gifted by Professor Oscar Marín, King’s College London). These strains were maintained at the Queensland Brain Institute, University of Queensland (QBI, UQ). Mice were housed in ventilated cages (Optimice, Animal Care Systems, CO, USA) and maintained at 21 ± 2°C ( 60 % humidity) with a 12-h light/dark cycle (lights on 0700h). Mice were housed in groups of two (with split dividers) per cage with *ad libitum* access to food and water. All experiments took place during the light phase.

*Fos^-2A-iCreER/+^; Ai9-tdTomato^f/+^* (TRAP2) mice were obtained by breeding *Fos^2A-iCreER/2A-iCreER^*mice (RRID: IMSR_JAX:030323; JAX stock #030323)(*15*) and *Ai9-tdTomato^f/f^* mice (RRID: IMSR_JAX:007909; JAX stock #007909)(*16*). *Pet-Cre^+/−^; Ai9-tdTomato^fl/+^*^;^*Erbb4^f/f^* mice were generated at the Institute of Neuroscience CSIC-UMH by breeding *Pet-Cre^+/−^* mice (Jackson Strain #:012712; RRID:IMSR_JAX:012712)(*17*), *Erbb4^f/f^* and *Ai9-tdTomato^fl/f^* (Ai9) mice(*16*) (generously gifted by Professor Beatriz Rico & Oscar Marín, King’s College London). Mice were housed under standard conditions on a 12 h light/dark cycle (lights on 0800h), with food and water available ad *libitum.* Mice were group-housed (eight mice per cage) in static microisolation cages with filter tops (Tecniplast), and were only single-housed when required by the experiment.

All experiments took place during the light phase. Females used in the study were in the Oestrus cycle, checked by visual inspection(*18*). All procedures were carried out in accordance with the Australian Code of Practice for the Care and Use of Animals for Scientific Purposes and the University of Queensland’s Institutional Biosafety Committee. Animal usage was approved by the University of Queensland Animal Experimentation Ethics Committee (AEC approval number 2022/AE000716) and the Ethics Committee of the Instituto de Neurociencias CSIC-UMH (2025-VSC-PEA-0082; Spain), according to both Spanish and European regulations.

### Activity-dependent neuronal labelling, imaging and quantification using TRAP2 animals

Adult TRAP2 mice were single-housed and habituated to the experimenter and to the transport to the behavioral room for three days. Mice received an intraperitoneal injection of tamoxifen (180 mg/kg) and were returned to their individual cages. One day later, mice were either exposed to the open field or stayed in the home cage. Three days later, animals were perfused, and their brains processed for analysis.

After image acquisition, whole-brain TRAP2 density was quantified according to the QUINT workflow(*19*). Segmentation was performed using Ilastik software10, and anatomical alignment to the Allen Mouse Brain Atlas (Common Coordinate Framework v3) was achieved with QuickNII and VisuAlign11,12. Cell counts for each brain region were obtained using Nutil13, based on Allen-defined anatomical areas. Brain heatmaps were generated using the N2U and MBH Shiny applications developed and previously employed by the Michael X. Handerson laboratory14,15. Pearson correlation coefficients between brain regions were calculated to construct cross-correlation matrices using the *cor* function in RStudio.

### scRNA-sequencing analysis

Publicly available dataset was downloaded from NCBI database (GEO accession: GSE144980) from Okaty et al., 2020 ((*20*), formed by 2,350 single *Pet1*^+^ cells from the dorsal raphe nucleus (DRN). QC was performed using the Seurat package, retaining for downstream analysis only cells with more than 4,500 detected genes; no additional gene-level filtering was applied, consistent with the threshold used in the original study. All 2,350 cells passed this filter. As detailed metadata were not available, we inferred biological sex using Seurat’s *AddModuleScore* function. Module scores were computed from well-established sex-linked genes (*Xist* for females; *Ddx3y, Kdm5d, Eif2s3y* and *Uty* for males), and cells were assigned a binary sex label (“female” and “male”) according to their higher score. We validated the segregation of both sexes using UMAP representation.

Sex-dependent gene expression was assessed with a single global comparison between female and male cells across all *Pet1^+^* neurons (not restricted by cluster), using Seurat’s *FindMarkers* function (*Wilcoxon* rank-sum test, the Seurat default; log fold-change threshold = 0, min.pct = 0.05). Results are reported as log_2_ fold changes, with positive values indicating overexpression in females and negative values indicating overexpression in males. Genes were classified as differentially expressed at FDR (Benjamini-Hochberg) < 0.05 and |log2FC| > 0.25; the full list of significant genes is provided in Table S1. Gene set enrichment analysis (GSEA) against Gene Ontology terms (Biological Process, Molecular Function and Cellular Component) was performed using *clusterProfiler::gseGO*, ranking genes by a signed score derived from their differential expression p-value (direction given by the sign of log2FC), with minGSSize = 10, maxGSSize = 500 and Benjamini-Hochberg-adjusted *p-values.* The percentage of *Erbb4*^+^ *Pet1*^+^ cells per sex was calculated as the proportion of cells with non-zero raw UMI counts for *Erbb4*.

To assess sex-dependent gene expression within a comparable cell population, we computed cell clustering following the original paper settings (PCs 1:5 and 8:50 combined, k. param = 20 in FindNeighbors; resolution = 0.9 in *FindClusters*) on the initial SCTransform-normalized embedding, used to visualize sex segregation (Figure 3A). For the analysis reported in Figure S3, a second, independent *SCTransform* normalization was performed regressing out the sex module scores (*sexF_1, sexM_1*) as covariates, followed by a newly computed PCA. Because this second embedding is derived from a distinct normalization and covariate structure, the specific principal components excluded in the first round do not necessarily correspond to the same axes of variation in the second; the full set of computed principal components (dims 1:50) was therefore retained for this clustering, following standard Seurat practice. This second clustering (resolution = 0.9) yielded 14 clusters, consistent with the original study. Molecular composition was evaluated using canonical markers from the original paper (*Tph2, Fev, Gad1, Gad2, Slc17a8* and *Met*), confirming an approximately similar cluster structure. Sex representation per cluster was assessed via *Xist/Ddx3y* UMAP projections and a stacked bar plot of the proportion of female and male cells per cluster.

### Behavioral Tests

Several behavioral assays were performed on male and female adult mice (P50+). Tests were performed in the order listed below, and mice were sex separated at analysis stages.

### Open Field

The open field recordings for the *Pet-Cre^+/−^; RCE-GFP* were conducted in a circular arena (39 cm diameter). Mice were habituated in the room with the open field at low light for 1 hour, then placed into the arena and recorded for 5 minutes from above as they freely roamed. Tracking analysis to determine parameters such as distance travelled and thigmotaxis was done via a MATLAB (Mathworks) code (*21*) modified for this specific open field arena. Briefly, the code masked the position of the mouse based on its binary contrast to the background arena. Distance was calculated by measuring the difference in the mask centroid position between each frame (Euclidean distance), then summated for the entire recording. Thigmotaxis was calculated by the ratio of time spent within the periphery of the arena compared to the centre, which was defined as an inner circle 5 cm away from the perimeter (*22*).

The open field used for TRAP2 and *Pet-Cre^+/−^; Ai9-tdTomato^fl/+^*^;^*Erbb4^f/f^* mice consisted of a square plastic chamber (50×50×50cm) illuminated at 70 lux. Mice were exposed to the test for 5 minutes, and their behavior was recorded with a Logitech C270 HD webcam. Analysis of the velocity, distance and center/border ratio was performed using EthoVision® XT software (Noldus, RRID:SCR_000441).

### Y-maze

The Y-maze spontaneous alternation task was performed in a transparent apparatus with 3 arms (50 x 8 cm each) under constant illumination of 70 lux. Mice were placed in the center of the maze and allowed to explore it for 8 minutes. Number of arm entries, velocity, and distance travelled were obtained using EthoVision® XT software. % Alternation was calculated as follows: % Alternation = total alternations / (total entries − 2) × 100.

### Sociability

Sociability and social novelty preference were evaluated using a three-chamber apparatus (50 × 25 × 25 cm) equipped with a cylinder in each side. The experiment consisted of three phases: habituation, a sociability phase (S1), and a social novelty phase (S2). During habituation, mice explored the empty arena for 5 min to confirm no baseline side preference. In the sociability phase, mice explored for 10 min with a novel mouse in one side chamber and an object in the other. In the social novelty phase, the object was replaced with a second novel mouse, and exploration was tracked for an additional 10 min. Chamber placements were counterbalanced across trials.

Exploration was defined as the time spent sniffing within a 15 cm zone around each cylinder, recorded using EthoVision® XT software. A discrimination index was calculated for each phase: for S1, by subtracting the time spent sniffing the object from the time spent sniffing the mouse subject; for S2, by subtracting the time spent sniffing the familiar subject from the time spent sniffing the unfamiliar subject.

### Light-Dark Box Test

Light-dark box tests (*23*) were performed in Imetronic Fear Cages (France). The dark box was in near darkness, and the light box was exposed to 400 lux light. Mice were placed into the dark box, with the door closed. The door was then opened, and the mouse allowed to roam freely between the rooms for 5 minutes. After the mouse completed the task and was removed from the chamber, the chamber was cleaned with a Virkon solution to remove excrement and odour cues. Time spent in each chamber within the 5 minutes was measured.

### Elevated Plus Maze

The elevated plus maze measures anxiety-like behavior in mice through the presentation of four crossing arms (62.5 x 5 cm), two of which are open, and two of which are enclosed with walls (height: 65 cm). Mice were habituated to the room for 30 minutes before being placed into the centre of the arena and allowed to roam freely for 5 minutes, under 200 lux. After the mouse completed the task and was removed from the chamber, the chamber was cleaned with an ethanol solution to remove excrement and odour cues. Analysis was done in Ethovision to determine the time spent in the centre of the arena and the open and closed arms. To assess anxiety-like behavior under less anxiogenic conditions, the EPM was also conducted under 100 lux illumination.

### Novelty suppressed feeding

The novelty suppressed feeding (NSF) test is an approach–avoidance task exploiting conflicting motivational states such as measuring the rodent’s exploratory drive against the innate drive to avoid danger (*24*). Animals were food-deprived for 16h prior to the test with continued access to water. The test arena was brightly lit (1100 lux) and consisted of a white rectangular box (50 cm L x 40 cm W X 20 cm H) with a 2 cm layer of bedding. A food pellet was attached to white filter paper over a petri dish and placed in the centre of the arena, level with the bedding. Mice were placed individually in the corner of the arena and were observed until they began to eat the pellet in the centre. The time until the first bite of food was recorded as the latency to eat. Mice were allowed a maximum of 10 minutes to approach and eat the food pellet. Following this, mice were returned to their home cage where they were allowed to eat a pre-weighed food pellet for a period of 5 minutes and the amount of food consumed was measured.

### Contextual fear conditioning

Contextual fear conditioning was performed using an acrylic chamber (17 × 17 × 25 cm) equipped with a shock-grid floor (Maze Engineers, USA), transparent walls, visual cues, and a 70% ethanol scent. On Day 1, mice explored the context for 3 min before receiving three foot-shocks (0.7 mA, 1 s duration, 30 s inter-shock interval). On Day 2, memory retrieval was assessed by returning mice to the chamber for 5 min without shocks. Freezing behavior was scored manually by an experimenter blinded to genotype and validated using EthoVision® XT software.

### Forced swim test

Mice were introduced into a water-filled transparent glass cylindrical tank (12 cm diameter, 23.5 cm height). Illumination was set to 56 lux, and the total experiment duration was 6 min. Behavior was recorded using a Logitech C270 HD webcam and manually analyzed *post-hoc* by an experimenter blinded to genotype and sex. Percentage of immobility was calculated to assess coping behavior.

### Tail suspension test

Mice were suspended by their tails using adhesive tape and positioned away from adjacent surfaces to prevent contact. A small plastic tube was placed over the base of the tail to prevent tail-climbing. Mouse behavior was recorded for 6 minutes under 56 lux illumination. Behavioral analysis was conducted manually by an experimenter blinded to genotype and sex, and the percentage of time spent immobile was calculated.

### Fiber photometry

*Surgery:* Mice were anaesthetised in an induction chamber with 5% isoflurane in oxygen and then mounted in a stereotaxic frame (Stoelting/RWD). Anaesthesia was maintained at 2% isoflurane throughout surgery. Prior to surgery, mice received a subcutaneous injection of buprenorphine for analgesia. A craniotomy was performed above the left basal lateral amygdala (BLA), and mice were injected unilaterally with an adeno-associated virus (AAV) encoding GRAB5-HT 1.0 (RRID; (*25*). AAV was delivered using either glass micropipettes (tip diameter 25–35 µm) connected to a Nanoject III microinjector (Drummond Scientific) or a NanoFil syringe (WPI) mounted on a UMP3 microsyringe injector (WPI). Viral vector was delivered at a rate of 75 nL/min using a Micro4 pump controller (WPI). The BLA was targeted relative to bregma at anteroposterior (AP) −1.5 mm, mediolateral (ML) +3.0 mm, and dorsoventral (DV) −4.75 mm, with a total injection volume of 350 nL. Following injection, the syringe was left in place for 10 min before being slowly retracted to allow viral diffusion and minimise backflow.

Following AAV injection, an optic fibre cannula (200 µm core diameter, numerical aperture 0.39, flat tip; RWD, Cat# R-FOC-BL200C-39NA) was implanted at the same AP and ML coordinates, at a depth of 4.70 mm. The implant was secured to the skull using Super-Bond dental cement (Sun Medical, Shiga, Japan). Mice were allowed to recover for 4 weeks before behavioural experiments. Following completion of the experiments, mice were perfused to verify correct optic fibre placement (Fig. 2C).

*Recordings:* To determine baseline serotonergic activity, animals were placed in a pseudo home cage (i.e. an identical Optimice cage with fresh bedding and no food and water). Animals were allowed to freely roam the pseudo home cage for 5 mins. Data was recorded using a Neurophotometrics FP3002 system (Neurophotometrics, San Diego, CA). Briefly, 470 and 415 nm LED light was bandpass filtered, reflected by a dichroic mirror, and focused onto a patch cord (Neurophotometrics, San Diego, CA) by a x20 objective lens. Alternating pulses of 470 and 415 nm light ( 50 µW) were delivered at 200 Hz, and photometry signals were acquired using a Bonsai workflow. Signals were analyzed using a custom-made Python script. To correct for photobleaching, isosbestic (415nm) signals were fit to a biexponential curve to model the baseline signal with photobleaching. The fitted biexponential curve was then scaled to match the 470nm signal using non-negative robust linear regression and then subtracted from the 470nm trace to correct for baseline drift. Traces were then Z scored.

*Experimental traces and analysis:* Z scored data were analysed by custom-built python scripts to create traces for each experiment. To find serotonergic events, a peak prominence threshold was set at 2.0. The number of peaks was then averaged over the 5 mins the animal was in the pseudo-home cage to determine event frequency. A nonparametric t-test (Mann-Whitney U test) was used to quantify 5-HT event frequency between sexes within our experimental cohort

### Immunohistochemistry

Mice (P60) were used for immunohistochemistry. Animals were transcardially perfused with cold 4% paraformaldehyde (PFA) in 0.1 M phosphate buffer (PB). Brains were postfixed for 2 h and subsequently cryoprotected in 15% and 30% sucrose in 0.1 M PB. Tissues were sectioned at 40 µm using a freezing microtome.

For immunohistochemistry, free-floating sections were blocked for 2 h at room temperature (RT) in PBS containing 0.3% Triton X-100 and 5% of normal goat serum (for 5-HT) or BSA (for ErbB4). Sections were then incubated overnight at 4°C with the corresponding primary antibodies in PBS containing 0.1% Triton X-100 and 1% of the respective blocking agent: rabbit anti-5-HT (1:2,500; Sigma-Aldrich Cat# S5545, RRID:AB_477522), rabbit anti-ErbB4 (1:300; 0618, produced by the Cary Lai laboratory) and goat anti-mCherry (1:500; Biorbyt, Cat# orb153320). After three 15-min washes in PBS, sections were incubated for 2 h at RT with the appropriate secondary antibodies and fluorescent detection reagents: donkey anti-goat Alexa Fluor 546 (1:1,000; Thermo Fisher Scientific Cat# A-11056, RRID:AB_2534103), biotinylated goat anti-rabbit antibody (1:1,000; Vector Laboratories Cat# BA-1000, RRID:AB_2313606) and Alexa Fluor 488– or 647-conjugated streptavidin (1:1,000; Molecular Probes Cat# S32354, RRID:AB_2315383 and Thermo Fisher Scientific Cat# S-21374, RRID:AB_2336066 respectively). After three 15-min washes in PBS, sections were counterstained with DAPI and mounted with Mowiol.

At the end of the photometry experiments, mice were perfused transcardially with 0.01M phosphate buffered saline (PBS) to eliminate blood and extraneous material, followed by 4% paraformaldehyde (PFA). Brains were postfixed for 36h in PFA. Tissues were sectioned at 100 µm using a Leica 1000S vibratome and kept in a cryoprotective ethylene glycol solution at –20°C until processed for immunofluorescence. Sections were first washed and permeabilized in PBS-Triton 0.25%, then non-specific binding sites were blocked by immersing the tissue in 10% normal donkey serum, 2% bovine serum albumin in PBS-Triton 0.25% during 2h. Tissues were then stained using the primary antibodies overnight: mouse anti-GFP (1:3000; Aves Lab). After 3x 15 min washes, we added anti-rabbit Alexa 488 (1:200; Life Technologies) secondary antibodies for 2h. After 3x 15 min washes slices were stained during 5 min with DAPI (5µM; Sigma), mounted on Superfrost Plus slides then coverslipped with Mowiol (Sigma) Imaging was conducted on a Zeiss 710 confocal fluorescent microscope (20x objective).

### Confocal Imaging and Analysis

Confocal images were acquired using a Leica SPE II upright microscope. A 20×/0.6 NA objective was used for cell distribution analysis, and a 40×/1.15 NA objective was used for fluorescence intensity measurements. Images used for fluorescence intensity analysis were acquired at 12-bit depth. Image analysis was performed using FIJI (ImageJ, RRID:SCR_002285)(*26*). The Cell Counter plugin was used for manual quantification of tdTomato-positive neurons and their colocalization with 5-HT or ErbB4. tdTomato-positive cell density was quantified semi-automatically and normalized to the analysed area. To verify expression of GRAB_5HT1.0_ and correct placement of the fibre, tiled images (10x objective) were acquired on a Zeiss LSM 710 inverted confocal microscope and automatically stitched together with Zen Black (Zeiss) software.

For ErbB4 fluorescence intensity analysis, tdTomato-positive cells were identified using a mask, which was then used to define the regions of interest for ErbB4 fluorescence measurements. Corrected total cell fluorescence (CTCF) was calculated as: integrated density − (cell area × mean background fluorescence). Individual cell CTCF values were used to generate cumulative frequency distributions.

For quantification of serotonergic neuropil in the BLA, the region of interest (ROI) was manually delineated in FIJI. A fixed background value and the same intensity threshold were applied to all images. The tdTomato-positive neuropil area was quantified and expressed as a percentage of the total BLA area.

Quantifications were performed in n ≥ 3 mice per condition using four sections per animal. For cell density and neuropil analyses, the mean value per animal was used for statistical analysis, whereas CTCF measurements were analysed at the single-cell level.

### Statistical analysis

Statistical analyses were performed using GraphPad Prism 9 (RRID: SCR_002798) or R software. Data are presented as mean ± SEM. Datasets were first tested for normality (Shapiro–Wilk test) and equal variance (Levene’s test). Normally distributed data were analyzed using parametric tests: unpaired two-tailed Student’s *t*-test, one-way ANOVA, or two-way ANOVA. Datasets that were not normally distributed were analyzed using non-parametric tests, such as the Mann–Whitney U test. Multiple comparisons were corrected using the Bonferroni method. Significance levels were set at p < 0.05 (*), p < 0.01 (**) or p < 0.001 (***).

### Custom-made codes

Custom built code to view and process raw photometry signals and calculate event frequency is available at https://github.com/elisecr/fiberphotometry-analysis.

Code for the scRNA-sequencing reanalysis (quality control, sex inference, clustering, differential expression, and gene set enrichment analysis) is available at https://github.com/aballesteros2605/scRNAseq-Serotonergic-5HT-cells.

## RESULTS

### Sex-differences in novelty-induced anxiety-like behavior

Sexual dimorphisms in anxiety-related behavior have been modelled in mice, although its reproducibility and characterization are variable depending on the parameters measured across different studies. This caveat is likely due to different behavioral paradigm and housing conditions which are difficult to reproducibly implement and parse out, especially when they have to be reflected in an averaging performance across large time epochs (>10min). Recent studies suggest that increasing the temporal resolution of behavioral analysis can reveal subtle sex differences that may be missed by conventional behavioral measures(*27*). To establish a robust and reproducible behavioral baseline upon which we could interrogate neurobiological mechanistic underlying sex dimorphisms beyond reproductive ones, we first systematically characterized behavioral response to anxiogenic stimuli using complementary paradigms that engage the conflict between exploration and threat avoidance. Our analysis employed a one-minute-long epoch dissection of mouse behavior to study the dynamics of adaptations to each anxiogenic context using sex as a biological variable.

During exposure to a novel open field arena that combines a rodent’s drive to explore with the anxiogenic challenge of an open, brightly lit space (70LUX), we observed that female mice exhibited significantly greater distance travelled and velocity compared to males (Fig. 1A). Females also displayed increased thigmotaxis, spending proportionally more time in the peripheral zone relative to males (Fig. 1B), indicating a more anxiogenic exploratory strategy under mild anxiogenic stimuli.I This initial phase of higher anxiety observed in females when compared to male littermates, was lost during subsequent 5-10 min phases of exploration of the novel open field, indicating adaptation to the novel open field in which sex-dimorphisms disappear (Fig. S1A-B). Critically, these behavioral differences were independently replicated across two laboratories under distinct housing conditions (Fig. S1A-C), confirming the robustness and reproducibility of the observed sex dimorphism during adaptation to a novel anxiogenic context (open field).

**Figure 1.**
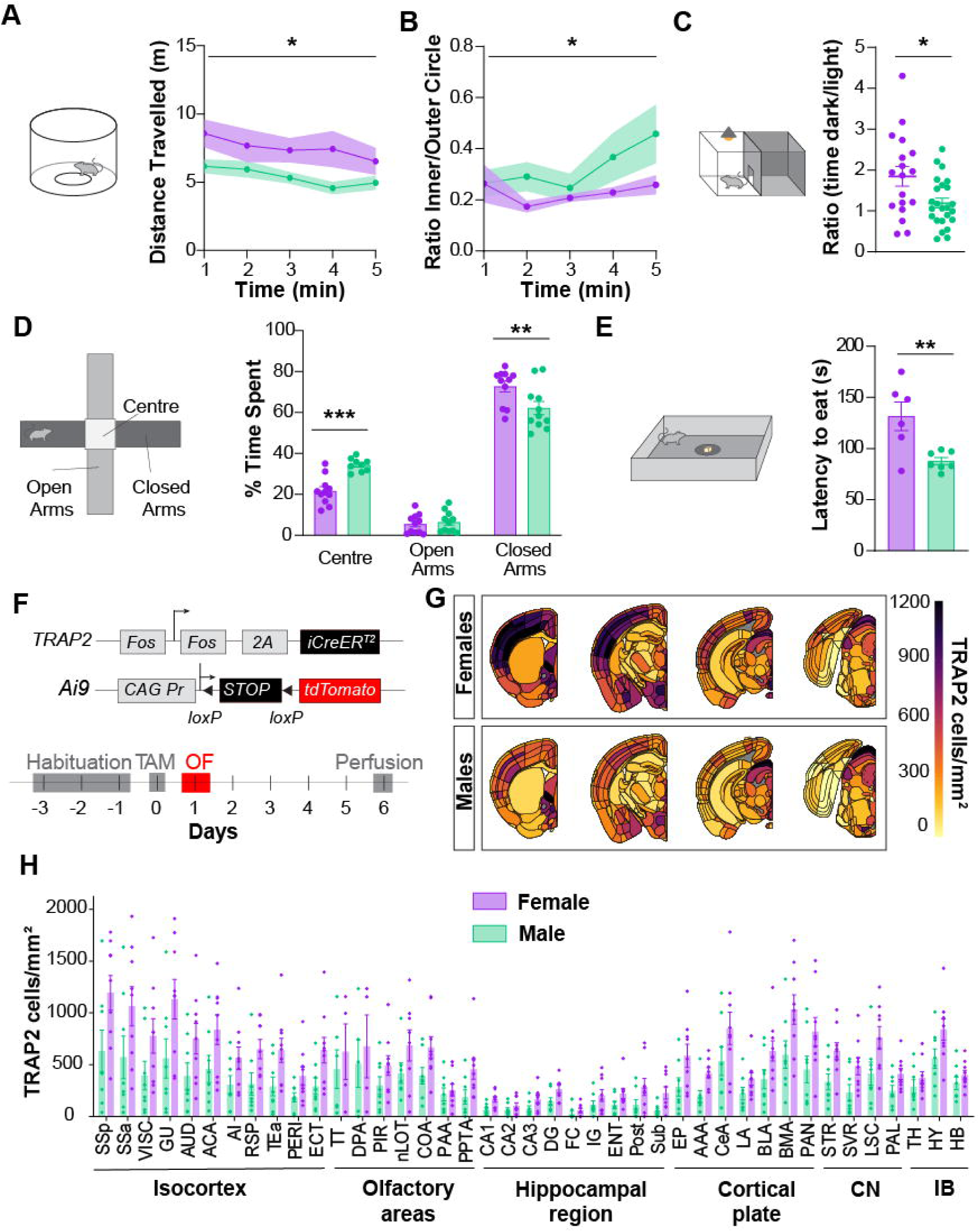
Sex-specific anxiety-like behaviour is correlated with divergent regional brain activation. **(A)** Total distance (m) travelled during initial exposure to the open field; n= 7 females, 9 males; p_sex_<0.0001 (2-way ANOVA). **(B)** Thigmotaxis test; n= 7 females, 9 males; p= 0.0361 (2-way ANOVA). **(C)** Dark/Light box test; n= 18 females, 24 males; p= 0.0115 (t-test). **(D)** Elevated Plus Maze test; n= 11 females, 11 males; P_Centre_— °0025, p_Open_>0.9999, p_closed_= 0.0003. **(E)** Force feeding test; n= 6 females, 7 males; p= 0.0079. **(F)** Experimental design with Fos TRAP2 mice. **(G)** Density map of TRAP2-positive cells. **(H)** Distribution of TRAP2-positive cells in the brain of female (n= 9) and male control mice (n= 8); p= 0.0039; CN: Cerebral nuclei; IB: Interbrain. See Fig.S2 for abbreviations. Data are presented as mean ± SEM. Statistical analysis and exact p values are in Table 1.

**Figure 2.**
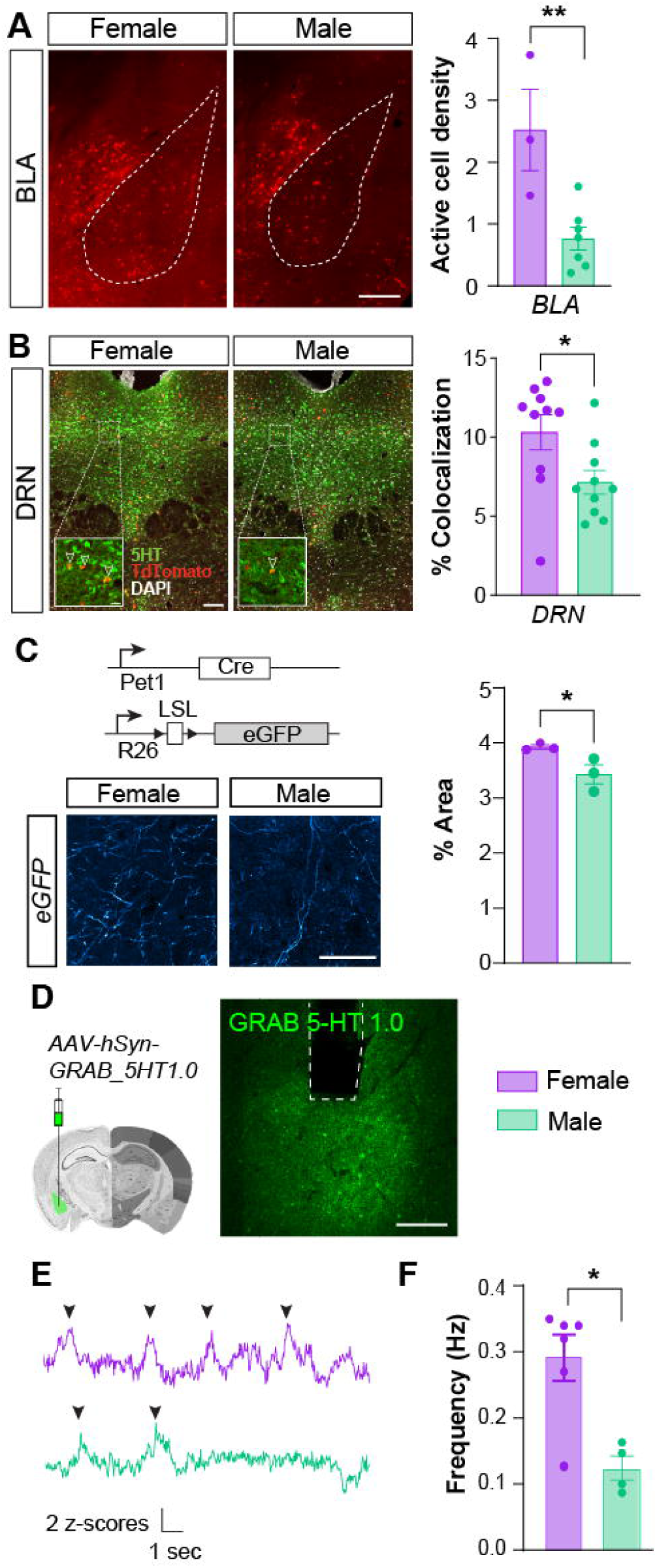
Enhanced serotonergic activity in females. **(A)** Active FOS+ cell density in the BLA in male and female mice. n= 3 females, 7 males; p= 0.0071, unpaired Student t-test. **(B)** Percent of active FOS+ cell density in serotonergic neurons of the DRN in male and female mice. n= 11 females, 10 males; p= 0.0167, unpaired Student t-test. **(C)** Fiber implantation in the BLA.Scale: 200 pm **(D)** Fiber recording traces in female and male mice. **(E)** Frequency of the GRAB 5HT signals; n= 6 females, 4 males; p= 0.0101, Mann-Whitney U Test. **(F) %** area occupied by neuropil in *Pet1-Cre; RCE-GFP* mice; n= 3 females, 3 males; p= 0.0470, unpaired Student t-test; Scale:50 pm. Data are presented as mean ± SEM. Exact p values are in Table 1.

To further assess sex-dimorphisms in anxiety-like behavior using an additional paradigm with increasing anxiogenic stimuli, we employed the light dark box test, which exploits the innate aversion of mice to strongly illuminated open spaces (400LUX). In line with a female-biased tendency to exhibit heightened response to aversive environments, female mice spent significantly more time in the dark compartment compared to males during the initial 5-min phase (Fig.1 C). Comparable sex-dimorphisms were found in another bona fide paradigm of anxiety-like behavior such as the elevated plus maze (Fig. 1D; S1D), where females spent significantly more time in the closed arms compared with males. To extend the same anxiety construct into a motivational and ethologically relevant domain (*i.e.* feeding behavior), rather than spatial exploration, we used a novelty suppressed feeding (NSF) test, which captures the suppression of an innate motivated behavior (feeding) in a novel anxiogenic context (*28*). We found that females displayed longer latency to eat than males in the arena (Fig.1E). However, we did not observe any differences in male vs female mice across a range of other behavioral paradigms including sociability (Fig. S1E-G), forced swim test (Fig. S1H), and tail suspension (Fig. S1I), to assess social behavior, stress-coping responses and behavioral despair (*29*, *30*), respectively. We next examined short-term spatial memory (through spontaneous alternation in the Y-maze) and fear memory (through contextual fear conditioning) and did not find any difference between males and females in their performance (Fig. S1J-L), indicating that males and females exhibited comparable fear learning and memory under our experimental conditions. Together the convergence of findings across independent behavioral paradigms suggests that females exhibit higher anxiety-like behavior during the initial phase of novel context exposure, accompanied by a distinct pattern of locomotor adaptation to the anxiogenic environment. Importantly, no sex differences in locomotor activity were observed under non– or milder anxiogenic conditions, such as the EPM at 100 Lux (Fig. S1L), indicating that the observed sex dimorphism in behavior is context-dependent rather that attributable to differences in locomotor capacity.

### Sex-differences in whole-brain recruitment upon exposure to a novel context are revealed by activity-dependent fluorescent labelling

We next examined functional differences in forebrain recruitment and neural circuit dynamics underlying sex-specific variation in novel context exploration. To capture behaviorally induced neural activation at the whole brain level, we employed activity-dependent labelling in *Fos-CreERT2; Ai9* mice. In line with prior studies (*31*), we established that tamoxifen administration 24h before a behaviorally relevant task, such as contextual fear conditioning, reliably induced activity-dependent fluorescent labelling at the whole brain level in regions known to be recruited by the contextual fear conditioning paradigm such as cortical (*e.g.* prefrontal cortex, anterior cingulate cortex) and hippocampal regions (data not shown). Before assessing sex differences, we tested whether tamoxifen metabolism differed between males and females. To this end, we induced activity-dependent labelling during their regular housing in the home cage, a familiar environment without behavioral novelty. Consistent with the absence of sex differences in behavioral parameters such as locomotor activity under these conditions, we observed no differences in activity-dependent labelling between males and females (Fig. S2A-C).

Next, we probed activity-dependent fluorescent labelling under novel context (open field) exposure (Fig. 1F). As previously observed in WT C57BL/6 mice, sex-differences in exploratory behavior during exposure to a novel context were also observed in *Fos-CreERT2; Ai9* mice. Specifically, female *Fos-CreERT2; Ai9* exhibited significantly increased distance travelled across the first minutes of exposure to an open field when compared to males (Fig.S2D). To monitor activity-dependent fluorescently labelled neural assemblies in an unsupervised and semi-automated fashion across the whole forebrain, we performed an atlas-based spatial quantification of tdTomato labelled cell density (mostly neurons) using QUINT workflow pipeline (*19*). Following manual alignment of brain slices to reference atlas, automated detection of fluorescently labelled neurons, our analysis determined that females exhibit significant higher level of activation of several forebrain regions when compared to males (Fig. 1G, H). Specifically, significant higher activation of hippocampal regions (CA1, CA3 and DG) as well as cortical and subcortical regions (STR).

### Sex-differences in serotonergic DRN recruitment and serotonergic wiring

Given the broad pattern of whole-brain activation observed when mice are placed in a novel anxiogenic context, we hypothesized that this global response is driven in part by the ascending serotonergic neuromodulatory system, whose extensive projections allow it to modulate global network excitability and behavior. Our prime target for investigating sex differences in whole-brain activation upon novel context exposure was the dorsal raphe nucleus (DRN) serotonergic subsystem. Serotonergic DRN neurons innervate virtually the entire forebrain and were shown to engage in emotionally salient stimuli (*32*) and to regulate a wide variety of emotional behaviors such as anxiety and social stress (*33*). To examine whether serotonergic neurons are differentially activated between male and female upon novel context exposure, we employed of activity-dependent fluorescent labelling upon novel context exposure. Serotonergic DRN neurons labelled by activity during open field exploration in TRAP2 (*Fos-CreERT2; Ai9)* mice were stained for 5-hydroxytryptamine (5-HT) and the degree of colocalization of TRAP2 and 5-HT labelled neurons was quantified in male and female mice. We first probed Fos+ cell density in the BLA and found a higher density of TdTomato+ cells in the female BLA (Fig. 2A). We also observed that, although the total number of 5-HT-stained DRN neurons was not significantly different between males and females (data not shown), the total number of activity-dependent (tdTomato+) labelled neurons in the DRN as well as the serotonergic DRN neurons tagged with activity dependent labelling (showing 5HT+ and tdTomato+ colocalization) was significantly increased in the female DRN, when compared to males (Fig. 2B). This data indicates a female-biased recruitment of serotonergic DRN neurons during open field exploration, suggesting that serotonergic DRN activation in a novel, anxiogenic context is higher in females than in males.

Since studies in both mice and humans have reported sex-dimorphisms in 5-HT receptor expression and signalling (*34*, *35*), we asked whether the sex-specific patterns of whole-brain activation evoked by novel context exposure might arise from differences in serotonergic innervation of downstream target regions. Mapping ascending 5-HT projections thus provides a crucial, mechanistically independent test of this hypothesis. Target-selective innervation could amplify or constrain the influence of DRN activity on specific postsynaptic regions, thereby shaping the distributed patterns of brain activation and behavioral outputs observed following novel context exposure. To address this, we used *Pet1-Cre* (*20*) mice bred with the GFP or Ai9 reporter mouse line that allowed us to visualize serotonergic axons (*36*). We quantified serotonergic axonal density in behaviorally relevant target sites such as the basolateral amygdala (BLA) and the hippocampus (*37*) using sex as a biological variable. Females exhibited significantly higher serotonergic axon density than males in both the BLA (Fig. 2C) and the border between stratum radiatum and lacunosum-moleculare (R-LM) of the hippocampus (S3 A, B). To further understand the basal functional readout of such organization, we performed *in vivo* fibre photometry in home cage 4 weeks following viral injection of GRAB 5-HT. We found higher release frequency in females compared to male littermates (Fig. 2 D-F), indicating sex-dependent differences in basal serotonergic activity. Overall, these findings indicate that sex-dependent differences in serotonergic circuitry extend beyond differential DRN recruitment to include differences in serotonergic innervation and basal 5-HT release dynamics, providing a potential circuit-level mechanism for sex differences in brain and behavioural responses to novel contexts.

### Sex-dependent molecular signatures in dorsal raphe serotonergic neurons

Building on our anatomical evidence of sex-dimorphic serotonergic projections, we sought to identify molecular factors acting within serotonergic neurons that could underlie these differences. In the adult mouse brain, serotonergic neurons display molecular heterogeneity, with distinct subtypes linked to specialized functions and region-specific projections (*38*). However, the molecular basis of sex-specific differences in serotonergic function has remained unexplored due to the absence of sex-based molecular profiling. To characterize the molecular basis of sex-dimorphisms in the DRN serotonergic subsystem, we reanalysed a publicly available single cell RNA sequencing (scRNAseq) dataset generated from recombinase-based genetic fate mapping of adult dorsal raphe serotonergic neurons using the *Pet1*-lineage (*20*) This dataset comprises 2,350 single cell transcriptomes with a mean of 7,521 genes detected per cell. As our principal goal was to characterize molecular pathways that differ between sexes in adult DR *Pet1*-lineage neurons, we first analysed and processed this scRNAseq dataset blind to the sex of the animal, as this information was not included in the originally published metadata. We inferred the biological sex of each profiled cell computationally, using Seurat’s *AddModuleScore* function to score each cell for a male gene module (*Ddx3y, Kdm5d, Eif2s3y* and *Uty*) and a female marker (*Xist*), and assigning each cell to the sex with the higher module score. This yielded 868 female-inferred cells (36.9%) and 1,482 male-inferred cells (63.1%); every cell received a binary label under this approach (Figure 3A).

To further characterised the different subtypes of *Pet1*^+^ cells withing the DR, we performed a clustering analysis of the neurons of the dataset. In the original study by Okaty et al. (2020) (*23*), PCA was performed on the scaled and centered expression values of the top 2,000 most variable genes. Principal components 6 and 7 were intentionally excluded from downstream clustering because their gene loadings were heavily weighted towards sex-specific transcripts. The published clustering therefore used PC1-5 and 8-50, yielding 14 transcriptionally distinct *Pet1*^+^ neuron subtypes. In order to explicitly assess sex-biased gene loading, we repeated the analysis using a second, independent *SCTransform* normalization that regressed out the female (*Xist*) and male (*Ddx3y, Kdm5d, Eif2s3y* and *Uty*) gene module scores as covariates, followed by a newly computed PCA. Because this second embedding reflects a distinct normalization and covariate structure from the first, we retained the full set of computed principal components (PC1-50, Seurat’s default) as input to *FindNeighbors*, rather than assuming the same PC6/7 exclusion criterion would apply to this independently derived embedding. Clustering was performed with the Seurat *FindClusters* function at a resolution of 0.9. Under these criteria, we reproduced a cluster structure closely matching the reported subtype marker genes *Tph2, Fev, Gad1, Gad2, Slc17a8* and *Met* (Figure S4A-B), indicating that our reanalysis successfully retained the expected expression profiles across the 14 clusters. Examination of X-linked gene *Xist* and Y-linked gene *Ddx3y* expression projected onto the UMAP embedding revealed that sex-linked transcripts were distributed broadly across clusters, consistent with sex-of-origin not being a primary driver of *Pet1*-neuron subtype identity (Figure S4C-D). Data indicate that, while sex-of-origin does not generally confound serotonergic subtype structure in the DR, some clusters represent transcriptionally distinct *Pet1* neuron subpopulations with a pronounced sex-biased cellular composition, suggesting that specific neuronal subtypes within the DR may be differentially represented or transcriptionally regulated in a sex-dependent manner (Figure S4E).

Differential expression analysis between sex-assigned neuronal populations performed on the full pseudobulk population of profiled Pet1+ neurons, rather than resolved separately within each transcriptional cluster, identified 1,779 genes significantly enriched in female neurons and 985 genes enriched in male neurons (FDR < 0.05, |log2FC| > 0.25) (Fig. 3B, Table S1). To identify the biological processes underlying these transcriptional differences, we performed Gene Set Enrichment Analysis (GSEA) on the ranked list of genes differentially expressed between male and female *Pet1*-lineage neurons, using Gene Ontology Biological Process, Cellular Component and Molecular Function gene sets. The top enriched GO terms revealed a pattern of sex-biased transcriptional programs: male-enriched genes were associated with mitochondrial and respiratory processes (cellular respiration, oxidative phosphorylation, respiratory chain complex), while female-enriched genes were associated with glycolysis and glucose catabolism, and with ribosomal subunit components (Figure S3F and Table S2). GO terms related to synaptic structure ranked near but did not cross de FDR < 0.05 threshold. Building on our anatomical evidence of sex-dimorphic serotonergic projections, we next asked whether any individual differentially expressed gene with an established role in axon guidance and circuit connectivity might help explain these projection differences. The Erb-B2 Receptor Tyrosine Kinase 4 (*Erbb4)*, encoding the receptor tyrosine kinase ErbB4 well characterized regulator of axon guidance (*39*) and synaptic connectivity(*40*), itself showed a statistically significant but quantitatively modest sex difference in expression (log2FC = –0.02, BH-FDR = 0.0009), reflecting the large number of cells analysed rather than a substantial change in expression level; percentage of *Erbb4*^+^ *Pet1*^+^ cells was 31.2% in females and 40.8% in males (Figure 3C).

Together, these results indicate that sex-dimorphism in gene-expression in adult DR *Pet1*-lineage neurons are broadly distributed across metabolic and trnaslsational programmes rather than concentrated in a single pathway, with only modest, sub-threshold trend toward male-biased synaptic gene expression. A point to consider in interpreting this analysis is that female– and male-derived cells were obtained from two separate sequencing captures rather than from multiple sex-balanced biological replicates. Thus, sex could be confounded with batch/capture in this dataset, and single-cell-level statistical tests could not be corrected by independent biological replication (see Limitations). These findings will therefore require validation in an independent, sex-balanced dataset before the identified pathways can be attributed to sex rather than technical variation.

### ErbB4 expression shapes sex-differences in serotonergic connectivity and influence the emergence of sex-dimorphisms in adaptive behavior

Through differential expression analysis of *Pet1*+ serotonergic neurons, we identified *Erbb4* (encoding the tyrosine kinase receptor ErbB4 also known as HER4, involved in axon guidance and circuit formation (*39*, *41*)) as a candidate gene within a sex-biased synaptic signalling pathway. To characterize ErbB4 expression at the protein level, we performed immunohistochemistry in adult male and female *Pet1^Cre/+^;tdTomato^RCL-flox/+^*mice (Figure 3E). ErbB4 immunoreactivity was detected in *Pet1*+ DRN neurons in both sexes. However, cumulative distribution analysis of corrected total cell fluorescence (CTCF) revealed significantly higher ErbB4 protein levels in *Pet1*+ DRN neurons from males compared to females (Figure 3E-F), suggesting sex-specific regulation of ErbB4 in DRN serotonergic neurons. To determine whether ErbB4 signalling contributes to sex-dimorphisms in adaptive behavior and sex-specific serotonergic circuit function, we generated a conditional knockout by crossing *Pet1^Cre/+^*mice with Erbb4*^flox^*and *tdTomato^RCL-flox^*animals (*Pet1^Cre/+^; Erbb4^flox/flox^;tdTomato^RCL-flox/+^*, henceforth Erbb4-cKO) (Figure 4D). First, we assessed exploratory behavior in the open field test. ErbB4-cKO males exhibited a marked increase in total distance travelled and mean velocity compared with male controls, whereas ErbB4-cKO females showed no significant change in distance or velocity during the first exposure to the open field (Figure 4A-B). This data indicates that ErbB4 in serotonergic circuits selectively constrains exploratory activity in males, consistent with its higher expression in male serotonergic neurons.

**Figure 3.**
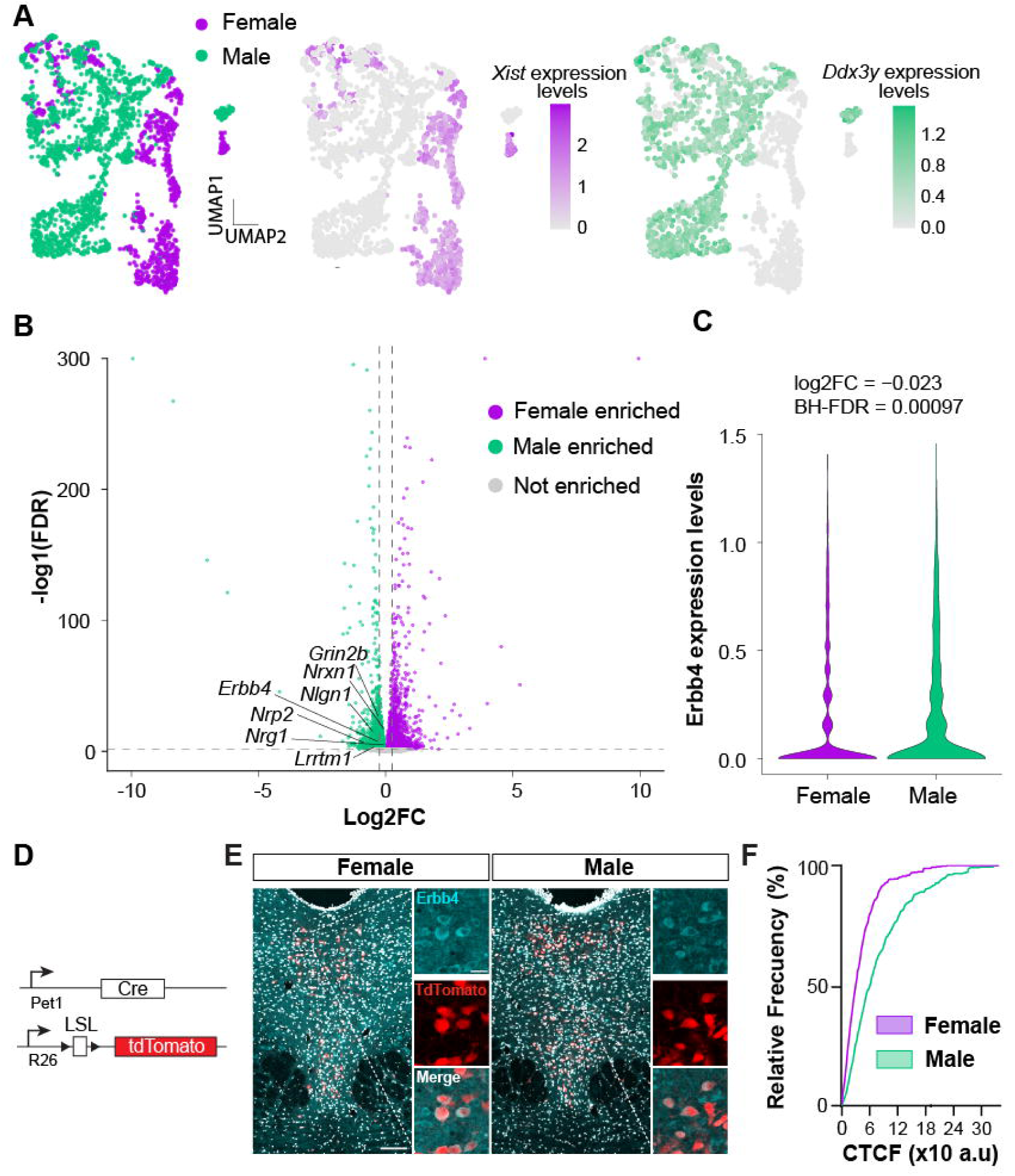
Sex-biased transcriptomic and protein-level profile of Pet1+ serotonergic neurons. (A) UMAP embedding of 2,350 single-cell transcriptomes from dorsal raphe Pet1-lineage neurons (Okaty et al., 2020; GSE144980). Left; cells colored by inferred sex identity (868 female-inferred, purple; 1,482 male-inferred, green). Middle/right: Xist and Ddx3y expression projected onto the same UMAP embedding. (B) Volcano plot of global differential gene expression between female– and male-inferred Pet1+ neurons (two-sided Wilcoxon rank-sum test, all cells per inferred sex, not restricted by cluster). Dashed lines mark the significance (BH-FDR < 0.05) and fold-change (|log2FC| > 0.25) thresholds used to call genes female-enriched (purple) or male-enriched (green); selected genes of interest are labeled. (C) Erbb4 expression in female-versus male-inferred Pet1+ neurons (same global comparison as in B; log2FC = –0.023, BH-FDR = 0.00097) and percentage of Erbb4+ Pet1+ cells per sex (31.2% female, 40.8% male). (D) Pet1Cre/+;tdTomato RCL-flox/+ gneetic strategy. (E) Representative confocal images of ErbB4 expression in Pet1-lienage neurons of the DRN. Scale barsJeft; 100 urn; upper right: 20 urn. (F) Cumulative frequency distribution of ErbB4 corrected total cell fluoresence (CTCF). n= 4 female and 3 male mice. Two way ANOVA with Kolmogorov-Smirnov post-hoc test (D=0.3588, p<0.0001); Interaction: P= 0.0003; row and column factor: < 0.0001. Exact p values reported in Tablel.

**Figure 4.**
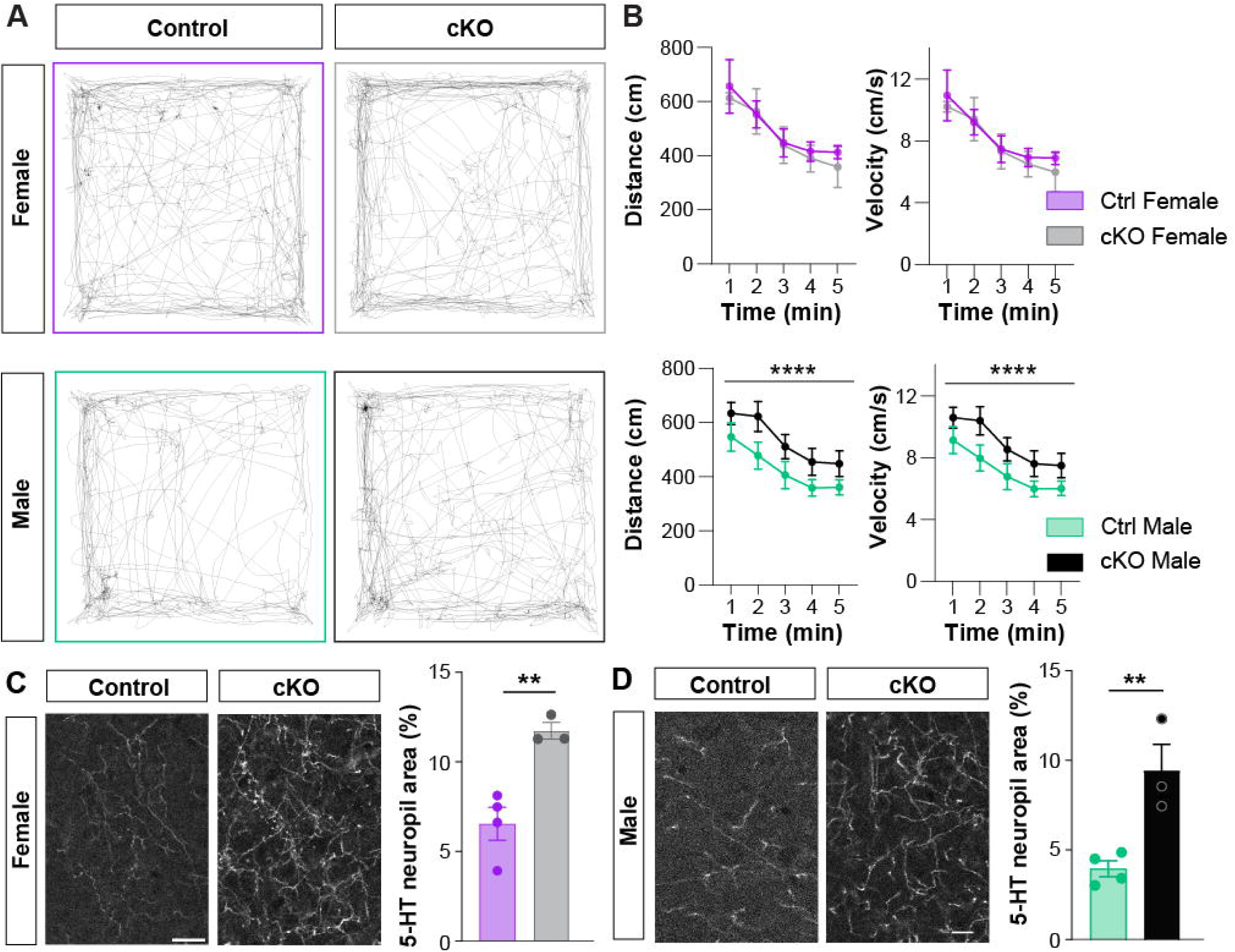
Sex-specific cellular and behavioral deficits in Pet1-positive serotonergic neurons of the DRN following ErbB4 ablation. (A) Open field recordings of controls (Pet1-Cre; Td-Tomato) and ErBB4 cKO (ErBB4 Pet1-Cre; Td-Tomato) female (top) and male (bottom) mice. (B) Distance and velocity of female (top; n= 7 controls, 5 ErBB4 cKO; p=0.0004 & p=0.0004, respectively) and male mice (Bottom; n= 12 controls, 11 ErBB4 cKO; p<0.0001). (C) Confocal images of Pet-Td-Tomato neuropil (left; scale: 20um) and 5HT neuropil area (%) in female mice; n= 4 control and 3 ErBB4 cKO female, p= 0.0039. (D) Confocal images of Pet-Td-Tomato neuropil (left; scale: 20um) and 5HT neuropil area (%) in male mice; 4 control and 3 *ErBB4* cKO male, p= 0.0027 Data are presented as mean ± SEM. Statistical analysis was performed using 2Way ANOVA Test. Exact p values are available in Table 1.

Finally, to assess whether ErbB4 loss alters serotonergic innervation of the BLA, we quantified Pet1-tdTomato neuropil area in this region. While a females showed a significant increase in BLA area occupied by Pet1-tdTomato neuropil when compared to males, ErbB4-cKO significantly increased 5-HT connectivity area in the BLA (Figure 4C-D), with no significant differences found between both sexes in ErbB4-cKOs. Together, these results indicate that ErbB4 expression in serotonergic neurons regulates serotonergic innervation in the BLA and shapes sex-dimorphic adaptive behavior.

## DISCUSSION

Sex as a biological variable is a major source of variability across a wide range of behaviors, including anxiety responses, ultimately contributing to the diversification of neural circuit function (*42*). However, the strong historical reliance on predominantly male animal models has left these neural circuit-level distinctions largely unexplored.

Leveraging recent advances in genetic and circuit-mapping technologies that enable precise delineation of sex-dimorphic serotonergic circuits in the mouse brain(*20*, *43*), we focused on the serotonergic system, a neuromodulatory system implicated in the regulation of these sex-dimorphic behaviors. Here, we identified a previously unrecognised temporal and context-dependent sex difference in behavioural adaptation to anxiogenic novelty, with females showing a stronger initial response that rapidly normalises with continued exposure. This behavioural dimorphism is accompanied by female-biased recruitment of dorsal raphe serotonergic neurons, greater serotonergic innervation of the hippocampus and basolateral amygdala, and increased basal 5-HT release, revealing sex-dependent differences in serotonergic circuit organisation and function. Integrating molecular and functional analyses, we identify ErbB4 as a candidate regulator of this dimorphism and show that its loss alters serotonergic BLA connectivity and selectively modifies adaptive behaviour in males.

### Structural basis of sex-specific behavioral adaptation

Here, our findings suggest that differences in the structural organization of serotonergic circuits contribute of sex-dependent behavioral differences, more specifically in the adaptation to a new and anxiogenic environment. We reveal sex-biased axonal occupancy of postsynaptic forebrain targets, including the basolateral amygdala. Using a novel context exploration paradigm in mice (*i.e.* an open, brightly lit arena), we discovered that a high temporal resolution of the analysis is critical for detecting such differences: Averaging behavior over extended periods masks sex differences. In contrast, analysing short time epochs during the initial exposure phase (*i.e.* first 5 minutes) revealed significant differences between males and females in distance travelled, velocity, and thigmotaxis (Fig. 1). These effects were consistently reproduced across two independent laboratories operating under distinct housing conditions, supporting the robustness of the phenotype. Similar observations in the elevated plus maze (Fig.1D) further suggest that this approach captures a generalizable feature of anxiety-related exploration.

From an ecological perspective, sex differences in anxiety-like behavior are likely to reflect adaptive strategies that evolved to optimize survival and reproductive fitness under different environmental pressures(*44*). In natural settings, males and females are exposed to partially distinct selective pressures, including territory exploration, resource acquisition, predator avoidance, reproductive competition, and parental investment. Subtle variations in risk assessment and exploratory behavior may therefore confer selective advantages by biasing behavioral responses toward strategies that are more beneficial for each sex within specific environmental contexts. Rather than representing maladaptive divergence, sex-biased anxiety responses may constitute different behavioral strategies by which each sex calibrates its response to threatening conditions. The behavioral differences observed in this study are likely influenced by the controlled conditions under which laboratory animals are maintained. Unlike wild populations, laboratory mice are raised in environments with reduced ecological complexity, limited predation pressure, stable food availability, and minimal social competition (*45*). Similar considerations have emerged from studies in other model organisms, including *Drosophila* (*46*) and *C. elegans* (*47*), where sexually dimorphic phenotypes often become more pronounced under ethologically relevant or environmentally challenging conditions. It is therefore plausible that sex-dependent behavioral adaptations are attenuated in standard laboratory settings and would be amplified in more naturalistic environments. Our findings additionally suggest that detection of sex differences critically depends on the intensity and temporal dynamics of anxiogenic stimulation. Serotonergic systems are organized into anatomically and functionally specialized subnetworks that can be recruited in a context-dependent manner (*43*). By focusing on the earliest phase of exposure to an anxiogenic environment, our paradigm may have preferentially captured rapid circuit-level adaptations that are otherwise diluted in conventional analyses over long time epochs.

Serotonin is involved in a broad range of behaviors extending beyond anxiety, including social interaction, sensory processing, learning, affective regulation, and cognitive flexibility (*49*). Despite growing evidence for sex differences in serotonergic function, including receptor signalling and behavioural responses to environmental challenges (*34*, *50*)whether the connectivity of DRN serotonergic neurons differs between males and females has remained largely unexplored. Although recent studies have revealed substantial heterogeneity in DRN serotonergic subsystems (*38*, *43*) and projection-specific functional responses(*32*), whether these circuit architectures differ between males and females remains largely unexplored (*32*). Our findings provide the first evidence that DRN serotonergic connectivity itself is sexually differentiated, suggesting that sex-dependent regulation of serotonin function may be established not only through cellular physiology but also through differences in circuit wiring. In addition, the higher frequency of spontaneous 5-HT transients in females may reflect a greater basal engagement of serotonergic signalling, potentially providing enhanced moment-to-moment regulation of internal and environmental states even in the absence of an overt challenge. The sex-biased serotonergic wiring and activity identified in this study may therefore support a wider repertoire of sexually dimorphic behavioral adaptations beyond defensive responses alone, according to environmental and physiological demands. Future investigations are warranted to confirm this assumption.

Another important question is whether the sex-biased serotonergic wiring identified here reflects the impact of circulating gonadal hormones or cell-intrinsic mechanisms that establish and maintain sex-biased circuit organization. Although gonadal hormones are well established modulators of serotonergic function at multiple levels (*51–54*), sex differences in anxiety-related behaviours have also been reported under conditions in which circulating sex hormones are low or experimentally controlled (*55*), suggesting that aspects of these behavioural differences may persist beyond specific gonadal hormone effects or reproductive stages. Sex differences have been reported in serotonin release and in the distribution of serotonin-related enzymes and in the subtype of serotonin receptor expression across different brain regions, indicating that serotonergic function is shaped by sex at multiple biological levels. In this context, the sex-biased connectivity identified here adds an additional layer of sex-dimorphism to the organization of the serotonergic system. Differences in axonal connectivity among target brain regions could provide a structural substrate for differential recruitment of downstream circuits, allowing serotonergic neurons, which axons collateralize prominently, to engage distinct sets of target regions in a sex-dependent manner. Such wiring organization may contribute to specificity of differential post-synaptic target regulation and might have been highly required for neuromodulatory systems such as serotonergic with a few thousands of cells regulating the entire forebrain while remaining sensitive to changes in circuit state induced by experience. Determining how this structural organisation interacts with hormonal state, developmental stage and experience will be an important direction for future studies.

### Molecular control of sex-dimorphic serotonergic circuits

The distinct molecular signatures observed in male and female serotonergic neurons suggest that these cells may rely on different biological strategies to support circuit function and behavioural adaptation. Notably, many of the enriched genes have fundamental roles in neuronal development, and the enrichment was not sex-exclusive but reflected a relative shift, with glycolytic metabolism and ribosomal/translational machinery more prominent in females, and mitochondrial/respiratory processes together with synaptic connectivity-related pathways more prominent in males. This suggests that sex differences in behavior and circuit architecture are reflected in subtle differences in molecular programs in the adult brain at the transcript level. Importantly, genes with similar RNA expression levels in males and females may nevertheless show different protein levels. It remains to be elucidated whether molecular programs show a more pronounced sex-bias at the protein level or during developmental periods. Thus, transcriptional similarity does not necessarily imply functional equivalence, and sex-specific developmental trajectories may emerge through differences in when and how shared molecular programs are deployed.

Investigating the molecular basis of sex-dimorphic circuit architecture, we identified the tyrosine kinase receptor *Erbb4* as a gene within a sex-differentially enriched pathway, the manipulation of which resulting in both large-scale remodeling of serotonergic projections (**Fig. 4** and (*36*)) and a loss of sex-specific behavioral adaptation to novel contexts. We propose two possible mechanisms through which *Erbb4* could regulate serotonergic circuit organization. First, ErbB4 may influence axonal guidance, sprouting or reorientation towards the amygdala (and CA1), potentially as a compensatory response to reduced innervation to other target regions, as previously described following manipulation of other guidance cues such as EphrinA5 (*56*). In previous studies, we showed that *Erbb4*-deficient serotonergic circuits fail to innervate the paraventricular thalamic nucleus (PVT) in both sexes. Thus, in line with the compensatory changes observed in serotonergic circuits upon ectopic expression of repulsive cues (EprinA5 (*56*)), the increased DRN-to-BLA innervation observed in Erbb4-deficient serotonergic circuits could represent a compensatory response to impaired DRN-to-PVT innervation, which requires *Erbb4* expression in both female and male. Second, ErbB4 may act through a synaptogenic mechanism, whereby presynaptic ErbB4 levels modulate synaptic connectivity (*40*, *41*), such that loss of Erbb4 function promotes increased axonal branching and sprouting due to the lack of synapse formation. Together, these possibilities point towards a role of Erbb4 in guiding axonal growth and target selection, with a potential dosage-dependent mechanism contributing to sex-biased circuit assembly, as hinted by the subtle but statistically significant different baseline levels of ErbB4 expression between males and females. Next investigations will have to decipher when and how Erbb4 in serotonergic cells impacts guidance molecules and/or synaptogenesis in longitudinal studies.

### Developmental underpinnings of the sex-dimorphic serotonergic circuits

Our findings support a model in which developmental organization of serotonergic circuitry contributes to sex-specific behavioral adaptation through selective and regionally biased circuit architecture, rather than arising solely from hormonal influences or receptor subtype-dependent mechanisms (*57*). Whether the structural differences in serotonergic connectivity are already evident at birth would need to be explored, to confirm whether these dimorphisms arise during early development rather than emerging solely through experience. This is consistent with prior work demonstrating that gonadal hormone signalling, particularly neonatal estradiol exposure, shapes serotonergic innervation patterns in a region-specific manner (*58*). How such hormonal cues interact with intrinsic developmental programs to generate sex-specific circuit architecture remains unresolved. Dissecting the relative contributions of gonadal hormones, sex chromosome complements, and their interactions will require targeted approaches, including hormone manipulation and sex chromosome models (*59*). Disruptions to serotonergic signalling during critical developmental windows are known to produce long-lasting consequences on adult emotional behavior, including heightened anxiety and altered stress responsivity (*e.g*., Pet1–dependent transcriptional switches that regulate postnatal maturation of serotonin neuron excitability; (*52*). Further dissecting how these mechanisms interact across developmental stages and within other neuromodulatory systems will be essential for understanding how early-life experiences and molecular programs converge to shape mature serotonergic function in both sexes. Advancing knowledge of sex-specific neural serotonergic circuit structure and function will not only uncover fundamental developmental processes but also will help elucidate the biological basis of neuropsychiatric disorders, including anxiety, ADHD and autism, to ultimately improve therapeutic outcomes for these conditions.

## ACKNOWLEDGMENTS

We thank Cari Lai (Indiana University, US) for their home-made anti-ErbB4 antibody (0618). ND thanks the Margaret Pemberton and Armstrong Foundations for their philanthropic support. The authors gratefully acknowledge the QBI Advanced Microscopy Facility for their support & assistance in this work. This work was supported by grant CNS2023-145536 [I.D.P] funded by MICIU/AEI/10.13039/501100011033 and by the European Union NextGenerationEU/PRTR.

## REFERENCES

1. J. Park, B. Moghaddam, Impact of anxiety on prefrontal cortex encoding of cognitive flexibility. Neuroscience 345, 193–202 (2017).

2. F. Meacham, C. T. Bergstrom, Adaptive behavior can produce maladaptive anxiety due to individual differences in experience. Evol., Med., Public Heal. 2016, 270–285 (2016).

3. C. A. Marcinkiewcz, C. M. Mazzone, G. D’Agostino, L. R. Halladay, J. A. Hardaway, J. F. DiBerto, M. Navarro, N. Burnham, C. Cristiano, C. E. Dorrier, G. J. Tipton, C. Ramakrishnan, T. Kozicz, K. Deisseroth, T. E. Thiele, Z. A. McElligott, A. Holmes, L. K. Heisler, T. L. Kash, Serotonin engages an anxiety and fear-promoting circuit in the extended amygdala. Nature 537, 97–101 (2016).

4. A. L. Garcia-Garcia, S. Canetta, J. M. Stujenske, N. S. Burghardt, M. S. Ansorge, A. Dranovsky, E. D. Leonardo, Serotonin inputs to the dorsal BNST modulate anxiety in a 5-HT1A receptor-dependent manner. Mol. Psychiatry 23, 1990–1997 (2018).

5. Y. Ohmura, K. F. Tanaka, T. Tsunematsu, A. Yamanaka, M. Yoshioka, Optogenetic activation of serotonergic neurons enhances anxiety-like behaviour in mice. Int. J. Neuropsychopharmacol. 17, 1777–1783 (2014).

6. X.-D. Yu, Y. Zhu, Q.-X. Sun, F. Deng, J. Wan, D. Zheng, W. Gong, S.-Z. Xie, C.-J. Shen, J.-Y. Fu, H. Huang, H.-Y. Lai, J. Jin, Y. Li, X.-M. Li, Distinct serotonergic pathways to the amygdala underlie separate behavioral features of anxiety. Nat. Neurosci. 25, 1651–1663 (2022).

7. L. Zhang, W. Ma, J. L. Barker, D. R. Rubinow, Sex differences in expression of serotonin receptors (subtypes 1A and 2A) in rat brain: a possible role of testosterone. Neuroscience 94, 251–259 (1999).

8. K. P. Cosgrove, C. M. Mazure, J. K. Staley, Evolving Knowledge of Sex Differences in Brain Structure, Function, and Chemistry. Biol. Psychiatry 62, 847–855 (2007).

9. S. Nishizawa, C. Benkelfat, S. N. Young, M. Leyton, S. Mzengeza, C. de Montigny, P. Blier, M. Diksic, Differences between males and females in rates of serotonin synthesis in human brain. Proc. Natl. Acad. Sci. 94, 5308–5313 (1997).

10. H. Jovanovic, J. Lundberg, P. Karlsson, Å. Cerin, T. Saijo, A. Varrone, C. Halldin, A.-L. Nordström, Sex differences in the serotonin 1A receptor and serotonin transporter binding in the human brain measured by PET. NeuroImage 39, 1408–1419 (2008).

11. C. Barth, A. Villringer, J. Sacher, Sex hormones affect neurotransmitters and shape the adult female brain during hormonal transition periods. Front. Neurosci. 9, 37 (2015).

12. K. N. Krolick, Q. Zhu, H. Shi, Effects of Estrogens on Central Nervous System Neurotransmission: Implications for Sex Differences in Mental Disorders. Prog. Mol. Biol. Transl. Sci. 160, 105–171 (2018).

13. D. A. Bangasser, A. Cuarenta, Sex differences in anxiety and depression: circuits and mechanisms. Nat. Rev. Neurosci. 22, 674–684 (2021).

14. J. D. Dougherty, N. Marrus, S. E. Maloney, B. Yip, S. Sandin, T. N. Turner, D. Selmanovic, K. L. Kroll, D. H. Gutmann, J. N. Constantino, L. A. Weiss, Can the “female protective effect” liability threshold model explain sex differences in autism spectrum disorder? Neuron 110, 3243–3262 (2022).

15. L. A. DeNardo, C. D. Liu, W. E. Allen, E. L. Adams, D. Friedmann, L. Fu, C. J. Guenthner, M. Tessier-Lavigne, L. Luo, Temporal evolution of cortical ensembles promoting remote memory retrieval. Nat. Neurosci. 22, 460–469 (2019).

16. L. Madisen, T. A. Zwingman, S. M. Sunkin, S. W. Oh, H. A. Zariwala, H. Gu, L. L. Ng, R. D. Palmiter, M. J. Hawrylycz, A. R. Jones, E. S. Lein, H. Zeng, A robust and high-throughput Cre reporting and characterization system for the whole mouse brain. Nat. Neurosci. 13, 133–140 (2010).

17. M. M. Scott, C. J. Wylie, J. K. Lerch, R. Murphy, K. Lobur, S. Herlitze, W. Jiang, R. A. Conlon, B. W. Strowbridge, E. S. Deneris, A genetic approach to access serotonin neurons for in vivo and in vitro studies. Proc. Natl. Acad. Sci. 102, 16472–16477 (2005).

18. S. L. Byers, M. V. Wiles, S. L. Dunn, R. A. Taft, Mouse Estrous Cycle Identification Tool and Images. PLoS ONE 7, e35538 (2012).

19. S. C. Yates, N. E. Groeneboom, C. Coello, S. F. Lichtenthaler, P.-H. Kuhn, H.-U. Demuth, M. Hartlage-Rübsamen, S. Roßner, T. Leergaard, A. Kreshuk, M. A. Puchades, J. G. Bjaalie, QUINT: Workflow for Quantification and Spatial Analysis of Features in Histological Images From Rodent Brain. *Front*. Neuroinformatics 13, 75 (2019).

20. B. W. Okaty, N. Sturrock, Y. E. Lozoya, Y. Chang, R. A. Senft, K. A. Lyon, O. V. Alekseyenko, S. M. Dymecki, A single-cell transcriptomic and anatomic atlas of mouse dorsal raphe Pet1 neurons. eLife 9, e55523 (2020).

21. C. Zhang, H. Li, R. Han, An open-source video tracking system for mouse locomotor activity analysis. BMC Res. Notes 13, 48 (2020).

22. M. L. Seibenhener, M. C. Wooten, Use of the Open Field Maze to Measure Locomotor and Anxiety-like Behavior in Mice. J. Vis. Exp., e52434 (2015).

23. M. Bourin, M. Hascoët, The mouse light/dark box test. Eur. J. Pharmacol. 463, 55– 65 (2003).

24. B. A. Samuels, R. Hen, Mood and Anxiety Related Phenotypes in Mice, Characterization Using Behavioral Tests, Volume II. Neuromethods, 107–121 (2011).

25. J. Wan, W. Peng, X. Li, T. Qian, K. Song, J. Zeng, F. Deng, S. Hao, J. Feng, P. Zhang, Y. Zhang, J. Zou, S. Pan, M. Shin, B. J. Venton, J. J. Zhu, M. Jing, M. Xu, Y. Li, A genetically encoded sensor for measuring serotonin dynamics. Nat. Neurosci. 24, 746–752 (2021).

26. J. Schindelin, I. Arganda-Carreras, E. Frise, V. Kaynig, M. Longair, T. Pietzsch, S. Preibisch, C. Rueden, S. Saalfeld, B. Schmid, J.-Y. Tinevez, D. J. White, V. Hartenstein, K. Eliceiri, P. Tomancak, A. Cardona, Fiji: an open-source platform for biological-image analysis. Nat. Methods 9, 676–682 (2012).

27. B. L. Smarr, A. D. Grant, I. Zucker, B. J. Prendergast, L. J. Kriegsfeld, Sex differences in variability across timescales in BALB/c mice. Biol. Sex Differ. 8, 7 (2017).

28. S. R. Bodnoff, B. Suranyi-Cadotte, D. H. Aitken, R. Quirion, M. J. Meaney, The effects of chronic antidepressant treatment in an animal model of anxiety. Psychopharmacology 95, 298–302 (1988).

29. V. Castagné, P. Moser, S. Roux, R. D. Porsolt, Rodent Models of Depression: Forced Swim and Tail Suspension Behavioral Despair Tests in Rats and Mice. Curr. Protoc. Neurosci. 55, 8.10A.1–8.10A.14 (2011).

30. A. Armario, The forced swim test: Historical, conceptual and methodological considerations and its relationship with individual behavioral traits. Neurosci. Biobehav. Rev. 128, 74–86 (2021).

31. X. Fan, J. Song, C. Ma, Y. Lv, F. Wang, L. Ma, X. Liu, Noradrenergic signaling mediates cortical early tagging and storage of remote memory. Nat. Commun. 13, 7623 (2022).

32. G. E. Paquelet, K. Carrion, C. O. Lacefield, P. Zhou, R. Hen, B. R. Miller, Single-cell activity and network properties of dorsal raphe nucleus serotonin neurons during emotionally salient behaviors. Neuron 110, 2664–2679.e8 (2022).

33. W.-J. Zou, Y.-L. Song, M.-Y. Wu, X.-T. Chen, Q.-L. You, Q. Yang, Z.-Y. Luo, L. Huang, Y. Kong, J. Feng, D.-X. Fang, X.-W. Li, J.-M. Yang, L. Mei, T.-M. Gao, A discrete serotonergic circuit regulates vulnerability to social stress. Nat. Commun. 11, 4218 (2020).

34. Y.-J. Luo, H. Bao, A. Crowther, Y.-D. Li, Z.-K. Chen, D. S. Tart, B. Asrican, L. Zhang, J. Song, Sex-specific expression of distinct serotonin receptors mediates stress vulnerability of adult hippocampal neural stem cells in mice. Cell Rep. 43, 114140 (2024).

35. E. L. Moses-Kolko, J. C. Price, N. Shah, S. Berga, S. M. Sereika, P. M. Fisher, R. Coleman, C. Becker, N. S. Mason, T. Loucks, C. C. Meltzer, Age, Sex, and Reproductive Hormone Effects on Brain Serotonin-1A and Serotonin-2A Receptor Binding in a Healthy Population. Neuropsychopharmacology 36, 2729–2740 (2011).

36. C. Barettino, Á. Ballesteros-Gonzalez, A. Aylón, X. Soler-Sanchis, L. Ortí, S. Díaz, I. Reillo, F. García-García, F. J. Iborra, C. Lai, N. Dehorter, X. Leinekugel, N. Flames, I. D. Pino, Developmental Disruption of Erbb4 in Pet1+ Neurons Impairs Serotonergic Sub-System Connectivity and Memory Formation. Front. Cell Dev. Biol. 9, 770458 (2021).

37. C. M. Teixeira, Z. B. Rosen, D. Suri, Q. Sun, M. Hersh, D. Sargin, I. Dincheva, A. A. Morgan, S. Spivack, A. C. Krok, T. Hirschfeld-Stoler, E. K. Lambe, S. A. Siegelbaum, M. S. Ansorge, Hippocampal 5-HT Input Regulates Memory Formation and Schaffer Collateral Excitation. Neuron 98, 992–1004.e4 (2018).

38. J. Ren, A. Isakova, D. Friedmann, J. Zeng, S. M. Grutzner, A. Pun, G. Q. Zhao, S. S. Kolluru, R. Wang, R. Lin, P. Li, A. Li, J. L. Raymond, Q. Luo, M. Luo, S. R. Quake, L. Luo, Single-cell transcriptomes and whole-brain projections of serotonin neurons in the mouse dorsal and median raphe nuclei. eLife 8, e49424 (2019).

39. G. López-Bendito, A. Cautinat, J. A. Sánchez, F. Bielle, N. Flames, A. N. Garratt, D. A. Talmage, L. W. Role, P. Charnay, O. Marín, S. Garel, Tangential Neuronal Migration Controls Axon Guidance: A Role for Neuregulin-1 in Thalamocortical Axon Navigation. Cell 125, 127–142 (2006).

40. I. del Pino, C. García-Frigola, N. Dehorter, J. R. Brotons-Mas, E. Alvarez-Salvado, M. Martínez de Lagrán, G. Ciceri, M. V. Gabaldón, D. Moratal, M. Dierssen, S. Canals, O. Marín, B. Rico, Erbb4 Deletion from Fast-Spiking Interneurons Causes Schizophrenia-like Phenotypes. Neuron 79, 1152–1168 (2013).

41. P. Fazzari, A. V. Paternain, M. Valiente, R. Pla, R. Luján, K. Lloyd, J. Lerma, O. Marín, B. Rico, Control of cortical GABA circuitry development by Nrg1 and ErbB4 signalling. Nature 464, 1376–1380 (2010).

42. T. J. Akiki, J. Jubeir, C. Bertrand, L. Tozzi, L. M. Williams, Neural circuit basis of pathological anxiety. Nat. Rev. Neurosci. 26, 5–22 (2025).

43. J. Ren, D. Friedmann, J. Xiong, C. D. Liu, B. R. Ferguson, T. Weerakkody, K. E. DeLoach, C. Ran, A. Pun, Y. Sun, B. Weissbourd, R. L. Neve, J. Huguenard, M. A. Horowitz, L. Luo, Anatomically Defined and Functionally Distinct Dorsal Raphe Serotonin Sub-systems. Cell 175, 472–487.e20 (2018).

44. D. C. Blanchard, Sex, defense, and risk assessment: Who could ask for anything more? Neurosci. Biobehav. Rev. 144, 104931 (2023).

45. S. H. Richter, J. P. Garner, H. Würbel, Environmental standardization: cure or cause of poor reproducibility in animal experiments? Nat. Methods 6, 257–261 (2009).

46. B. J. Dickson, Wired for Sex: The Neurobiology of Drosophila Mating Decisions. Science 322, 904–909 (2008).

47. Z. A. Hilbert, D. H. Kim, Sexually dimorphic control of gene expression in sensory neurons regulates decision-making behavior in C. elegans. eLife 6, e21166 (2017).

48. A. F. Scheyer, O. J. Manzoni, The Map and the Territory: Mapping the Territory Regulated by Serotonergic Signaling at Striatal Projection Neurons. Neuron 98, 679– 680 (2018).

49. P. Salvan, M. Fonseca, A. M. Winkler, A. Beauchamp, J. P. Lerch, H. Johansen-Berg, Serotonin regulation of behavior via large-scale neuromodulation of serotonin receptor networks. Nat. Neurosci. 26, 53–63 (2023).

50. P. Pavlidi, N. Kokras, C. Dalla, Sex Differences in Brain Function and Dysfunction. Curr. Top. Behav. Neurosci. 62, 103–132 (2022).

51. C. L. Bethea, M. Pecins-Thompson, W. E. Schutzer, C. Gundlah, Z. N. Lu, Ovarian steroids and serotonin neural function. Mol. Neurobiol. 18, 87–123 (1998).

52. O. T. Hernández-Hernández, L. Martínez-Mota, J. J. Herrera-Pérez, G. Jiménez-Rubio, Role of Estradiol in the Expression of Genes Involved in Serotonin Neurotransmission: Implications for Female Depression. Curr. Neuropharmacol. 17, 459–471 (2019).

53. D. R. Rubinow, P. J. Schmidt, C. A. Roca, Estrogen–serotonin interactions: implications for affective regulation. Biol. Psychiatry 44, 839–850 (1998).

54. G. Fink, B. Sumner, R. Rosie, H. Wilson, J. McQueen, Androgen actions on central serotonin neurotransmission: relevance for mood, mental state and memory. Behav. Brain Res. 105, 53–68 (1999).

55. J. R. Boivin, D. J. Piekarski, J. K. Wahlberg, L. Wilbrecht, Age, sex, and gonadal hormones differently influence anxiety– and depression-related behavior during puberty in mice. Psychoneuroendocrinology 85, 78–87 (2017).

56. T. Teng, A. Gaillard, A. Muzerelle, P. Gaspar, EphrinA5 Signaling Is Required for the Distinctive Targeting of Raphe Serotonin Neurons in the Forebrain. eNeuro 4, ENEURO.0327-16.2017 (2017).

57. P. Gaspar, O. Cases, L. Maroteaux, The developmental role of serotonin: news from mouse molecular genetics. Nat. Rev. Neurosci. 4, 1002–1012 (2003).

58. A. M. K. Madden, A. T. Paul, R. A. Pritchard, R. Michel, S. L. Zup, Serotonin promotes feminization of the sexually dimorphic nucleus of the preoptic area, but not the calbindin cell group. Dev. Neurobiol. 76, 1241–1253 (2016).

59. G. J. D. Vries, E. F. Rissman, R. B. Simerly, L.-Y. Yang, E. M. Scordalakes, C. J. Auger, A. Swain, R. Lovell-Badge, P. S. Burgoyne, A. P. Arnold, A Model System for Study of Sex Chromosome Effects on Sexually Dimorphic Neural and Behavioral Traits. J. Neurosci. 22, 9005–9014 (2002).

